# Identification and genetic characterization of Jingmen tick virus from ticks sampled in select regions of Kenya;2022-2024

**DOI:** 10.1101/2025.07.24.666544

**Authors:** Lydia Mwasi, Samoel Khamadi, Wallace Bulimo, Johnson Kinyua, Santos Yalwala, Luis Pow Sang, Haynes Robert, Gerald G. Kellar, John Eads, Fredrick Eyase

## Abstract

Jingmen tick virus (JMTV), an emerging segmented RNA virus classified as an ungrouped flavivirus, poses a growing public health concern globally. Known for its association with febrile illnesses and wide host range, JMTV has been detected in Rhipicephalus, Hyalomma, and Amblyomma ticks collected from cattle, goats, sheep, camels, and chickens in pastoral regions of Kenya, including Baringo, Mandera, Malindi, Lamu, Mombasa, Wajir, Isiolo, and West Pokot. Using viral metagenomics next-generation sequencing, this study analysed adult ticks (n=1547, 72 pools). A total of 53% (38/72) pools were positive for at least one viral pathogen, with JMTV detected in 87% (33/38) of these pools across all study sites.

Phylogenetic analyses revealed evidence of distinct Kenyan JMTV strains, with sequence segments from Malindi and Wajir clustering uniquely in their own clade; suggesting potential localised evolutionary pressures. Time calibrated phylogeny for the segment 1(RdRp) suggested varied ancestral origins and evolutionary relationships for the JMTV strains. MEME, BUSTED and FUBAR methods implemented in the Data-Monkey, unanimously identified codon 290 in segment 1 and 30 in segment 4 to be undergoing episodic positive selection. Recombination analysis performed using the RDP4 recombination detection tool indicated a recombination event in segment 2 of the Lamu JMTV strain that was confirmed by seven detection methods and visualised in BootScan.

These findings suggest that Kenyan JMTV strains are undergoing positive selection, potentially driven by unique ecological and host factors. Segmented genome evidence of recombination highlights the increasing virus’s potential for antigenic diversity. Host diversity and virus phylogenetic patterns underscore the zoonotic potential and its capacity for regional spread, emphasizing the critical need for enhanced vector surveillance. Temporal and ecological drivers like seasonal tick activity and livestock movement warrant investigation to elucidate JMTV transmission dynamics. Prioritizing tick-borne virus surveillance in Kenya will strengthen public health strategies and mitigates emerging viral risks.

## Introduction

Emerging and re-emerging pathogens are a great public health concern due to their potential to cause outbreaks [1]. Jingmen viruses have recently been identified and taxonomically described as unclassified flaviviruses [1, 2]. The first Jingmen virus was isolated in China from a *Rhipicephalus microplus* tick in 2010 [2]. The virus has two phylogenetic clades; one clade covers viruses from ticks, mosquitoes, humans, cattle, monkeys, bats, rodents, sheep, and tortoises and has been observed to cause febrile illness and flu-like symptoms in humans [2]. The second clade has been isolated from insect species, crustaceans, plants, and fungi [2]. Jingmen tick virus (JMTV) has a wide host range and viral transmission among ticks is efficient through “cofeeding”; it is termed as a true arbovirus, with a cycle involving an arthropod vector and a vertebrate host [3]. The fact that the virus has a wide host range and is distributed across different regions, makes it an emerging pandemic capable virus.

Unlike flaviviruses, the Jingmen virus is segmented with 4 to 5 segments; segment 1 of the Jingmen virus encodes for Non-Structural Protein 1 (NSP1), and resembles the flavivirus NS5-like protein that includes an RNA-dependent RNA polymerase (RdRp) and methyltransferase [4]. Segment 2 encodes for Viral Protein 1(VP1); Segment 3 encodes for Non-Structural Protein 2 (NSP2), which also shares similarities with the NS2B/NS3 flavivirus protein complex that encompasses transmembrane regions, serine protease and helicase domains, while Segment 4 encodes the Viral Protein 2-3 (VP2-3) [4]. Segments 2 and 4 are suggested to be structural proteins that constitute the envelope, capsid, and membrane proteins, respectively [4]. In addition, an open reading frame (ORF) overlapping with VP1 coding sequence at the beginning of segment 2 (VP4) is documented to encode for a small membrane protein [5]. The possible ancestral origin of non-structural proteins of segment 1 (NSP1) and segment 3 (NSP2) of the JMTV from flaviviruses suggest a unique evolutionary link between segmented Jingmen viruses and the non-segmented flaviviruses [6]. The overall length of the JMTV genome is 11,401 nucleotides, similar to that of flaviviruses [4].

In Kenya, JMTV has previously been isolated from Rhipicephalus, Hyalomma and Amblyomma tick species from goats, sheep, tortoise and cattle in Kajiado and Baringo pastoral regions [6]. Although there is evidence of circulation of the virus in these two pastoral regions in Kenya, there is no surveillance data of the virus in other pastoral regions of Kenya. This study addresses this gap by conducting a cross sectional study across select pastoral regions including Baringo, Mandera, Kilifi, Lamu, Mombasa, Wajir, Isiolo, and West Pokot counties. Baringo county previously reported Jingmen virus circulation [6], the Mandera county borders Ethiopia to the North, Wajir county borders Somalia to the West, Lamu county borders Somalia to the North and West Pokot county borders Uganda to the East; this trends towards surveilling for possible cross-border importation of vectors and consequently pathogen transmission due to transboundary movement of livestock. The pastoral nature of the study counties offers valuable insights into the prevalence and diversity of Jingmen tick virus in Kenya, highlighting its spread across the country and possible transmission to humans.

Unbiased metagenomic next generation sequencing (mUNGS) enables the identification of total microbiomes, making the identification of novel and emerging pathogens possible. In the present study Jingmen tick virus was identified and characterized, using this method.

## Materials and methods

### Ethical approval

The study was approved by the Kenya Medical Research Institute (KEMRI) Scientific and Ethics Review Unit (SERU) under protocol number KEMRI/SERU/CVR/4845 and license number 32115 by the National Commission for Science, Technology & Innovation (NACOSTI). The study was also submitted for approval to the Walter Reed Army Institute of Research (WRAIR) Human Subjects Protection Branch (HSPB) and Institutional Review Board (IRB) under package WRAIR # 3192 and was exempted from requiring any further approval as per WRAIR policy #25 because no samples, data or information was being collected from human subjects. Consent was sought from farmers and pastoralists using a written informed consent form.

### Sampling and pooling

Ticks were collected from cattle, sheep, goats, and camels from various pastoral counties including West Pokot, Isiolo, Baringo, Mandera, Lamu, Wajir and Kilifi (Fig 1). In Lamu, sample collection was carried outduring the month of October in Lamu, with 37mm precipitation and temperatures of 86-88°F. In Mandera, collection was done in June where temperatures of 67.5 - 84°F, precipitation of 1mm and humidity of 50% were experienced. In Isiolo, collection was done in March with temperatures of 74-84°F and precipitation of 43mm In Kilifi, collection was done in November with temperatures of 73.4-87.8°F, precipitation of 112.2mm and humidity of 81%. In Baringo, collection was done in November, with temperatures of 75-86°F and 58mm of precipitation. In West Pokot, collection was done in March, with temperatures of 54-79°F, precipitation of 67.5mm and humidity of 61%. Lastly in Wajir, collection was done in February, with temperatures of 75-97°F, precipitation of 2.1mm and humidity of 53%. The collected ticks were placed in 15 mL centrifuge tubes (Corning Inc, Corning, NY, USA) and transported to the KEMRI/ Walter Reed Army Institute of Research-Africa (WRAIR-A) laboratories on dry ice. The ticks were morphologically identified using taxonomical keys [7] and pooled in groups of 1 to 8 ticks per species and site of collection and stored at −81°C prior to laboratory methods [6].

**Fig 1.**
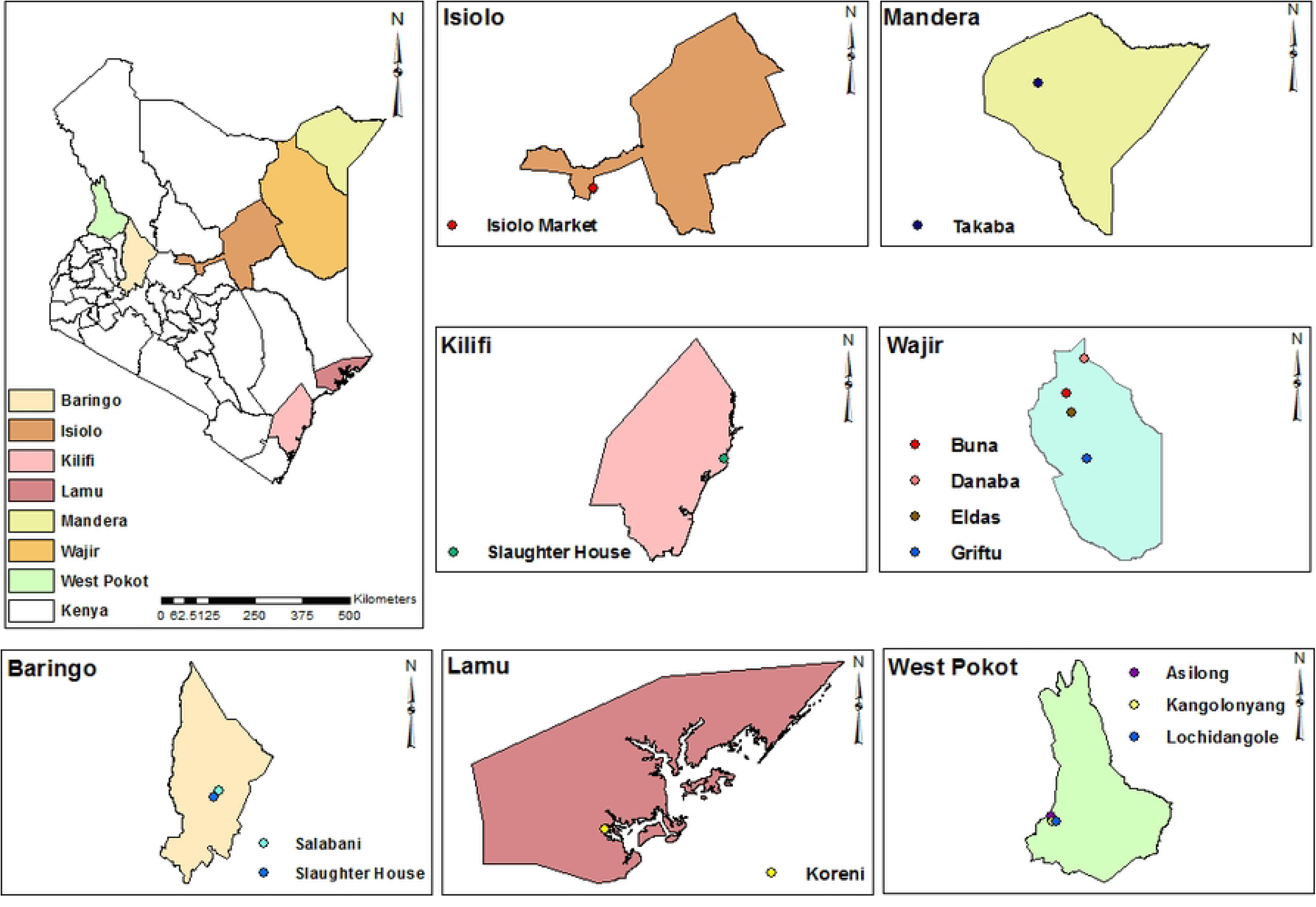
Tick sampling sites. This map was generated using ArcGIS version 10.2.2 for Desktop (Advanced License).

### Tick homogenization

Frozen individual pools of ≤10 ticks were homogenized with 0.7 ml dry volume 2.0 mm Zymo Bashing Beads (ZR S6003) in a mini-beadbeater-24 (Fisherbrand™) for 1 minute, with a mid-way pause of 10 seconds. One mL homogenizing media containing minimum essential medium, with Earle’s salts and reduced sodium hydrogen carbonate (NaHCO3), 15% fetal bovine serum (Gibco), 2% L-glutamine (Sigma Aldrich) and antimycotic/antibiotic solution (Sigma Aldrich) was added. The homogenates were vortexed and clarified through 10 minutes of 10,000 X g centrifugation (Eppendorf 5430-R) and filtered through a 0.22µm syringe filter (MilliporeSigma™, USA) to remove tissue debris and reduce quantities of bacteria and other impurities [8,9]. The homogenates were stored at −80°C prior to RNA extraction and metagenomics.

### RNA extraction and metagenomics

Total RNA was extracted from 100 µL of tick homogenates using the Zymo Quick-RNA™ Kit (USA), following the manufacturer’s instructions that included a DNase digestion step to digest free nucleic acid. The remaining unfiltered homogenates were stored at −80°C for viral isolation.

A sequencing library was prepared using the Illumina Ribo-Zero Plus Ligation kit (Illumina, USA) following the manufacturer’s instructions. RNA was quantified on a Qubit machine before library preparation using the RNA High Sensitivity (HS) kit (Invitrogen™) following ribosomal RNA depletion and DNA High Sensitivity (HS) kit (Invitrogen™) following cDNA synthesis. The 2100 Bioanalyzer (Agilent™) was used to quantify the final library before pooling and loading on the Illumina Miseq platform for sequencing with 300 *2 cycles.

### Sequence analysis

Sequence analysis was performed in Terra.bio [10] using the Theiameta_illumina_Pe_version PHB 2.1.0 workflow, where read quality trimming and adapter trimming was performed using the Trimmomatic tool v0.39 [11]. Read quantification was performed using fastq-Scan v0.4.4 [12], and taxonomic classification was performed in Kraken2 v2.1.2 [13]. De novo metagenomic assembly was performed using metaSPAdes v3.12.0 [14]; assembly alignment and contig filtering was done by SAMtools v1.17 [15] and Minimap2 v2.22-r1101[16] using a reference genome from NCBI. Comprehensive microbial variant detection and genome assembly improvement was performed by in Pilon v1.24 [17] and assembly quality assessment performed by the quality assessment Tool (QUAST) v5.0.2 [18] to give FASTA format contigs.

### Phylogenetic analyses

Maximum likelihood (ML) phylogenetic trees were estimated using Jones-Taylor-Thornton (JTT) model based on comparison of translated amino acid sequences of the four JMTV segments in MEGA 7 v7.0.26 with 1000 bootstrap replications nodal support [19,20], viewed using Figtree v1.4.4 [21] and midpoint rooted. Additionally, a phylogenetic time calibrated tree was generated for the study segment 1 genomes in BEAST v1.10. Briefly, 85 JMTV sequences were obtained from GenBank by country of origin and year of sample collection for inclusion in the calibrated tree analysis. Sequences were aligned using MAFFT software v7.310 and editing was performed in BioEdit software v7.7.1. BEAST was used to estimate the evolutionary rate in segment 1 (RdRp) by the relaxed uncorrelated lognormal molecular clock model. The maximum clade credibility (MCC) tree was run with a chain length of 250,000,000 replications; confirmation of convergence of all parameters in Tracer v1.7.2 (effective sampling size (ESS)> 200). The final MCC tree was generated by TreeAnnotator with a Burn-in of 10% of the total chain length and the tree was visualized by Figtree v1.4.4.

### Natural selection and recombination analyses

A mixed-effects maximum likelihood approach for individual sites under episodic positive selection was performed using the Mixed Effects Model of Evolution (MEME) analysis in Data monkey [22,23]. The results were confirmed using the Fast, Unconstrained Bayesian AppRoximation (FUBAR) [24] and the Branch-Site Unrestricted Statistical Test for Episodic Diversification (BUSTED) approaches in Data monkey [25].

Recombination analysis was performed using seven methods RDP, GENECONV, BOOTSCAN, MaxChi, Chimaera, SiScan and 3Seq implemented in RDP4 to identify and confirm potential recombination [26,27]. At least four of the seven methods had to detect recombination to consider a recombination event as significant and accurate; p-value cut off of 0.05 [28]. SimPlotv3.5.1 was used to generate recombination plot maps for visualisation and validation of recombination events detected by RDP4[29].

## Results

### Tick collection and morphological identification

A total of 1,629 ticks were collected from the study counties as follows: Isiolo (Market n=334); Lamu (Koreni n=25); Kilifi (Malindi, Slaughter house n=138); Mandera (Takaba n=147); Baringo (Salabani n=300,Slaughter house n=116); Wajir (Griftu n=102,Danaba n=39,Eldas n=43 and Buna n=2) and West Pokot (Lochidangole n=48,Kangolonyang n=9, Asilong n=88) (Table 1).The ticks collected in this study belonged to 4 genera and 14 species (Fig 2): *Amblyomma gemma* (13.5%), *Amblyomma lepidium*(6.3%), *Amblyomma variegatum*(7.7%), *Argas persicus* (18.4%), *Hyalomma albiparmatum* (2.3%), *Hyalomma anatolicum anatolicum* (0.4%),*Hyalomma dromedarii* (4.1%), *Hyalomma impeltatum* (0.4%), *Hyalomma marginatum rufipes*(17.4%), *Hyalomma truncatum* (7.9%), *Rhipicephalus appendiculatus* (4.2%), *Rhipicephalus boophilus microplus* (9.8%), *Rhipicephalus evertsi evertsi* (3.3%), and *Rhipicephalus pulchellus* (4.4%).

**Fig 2.**
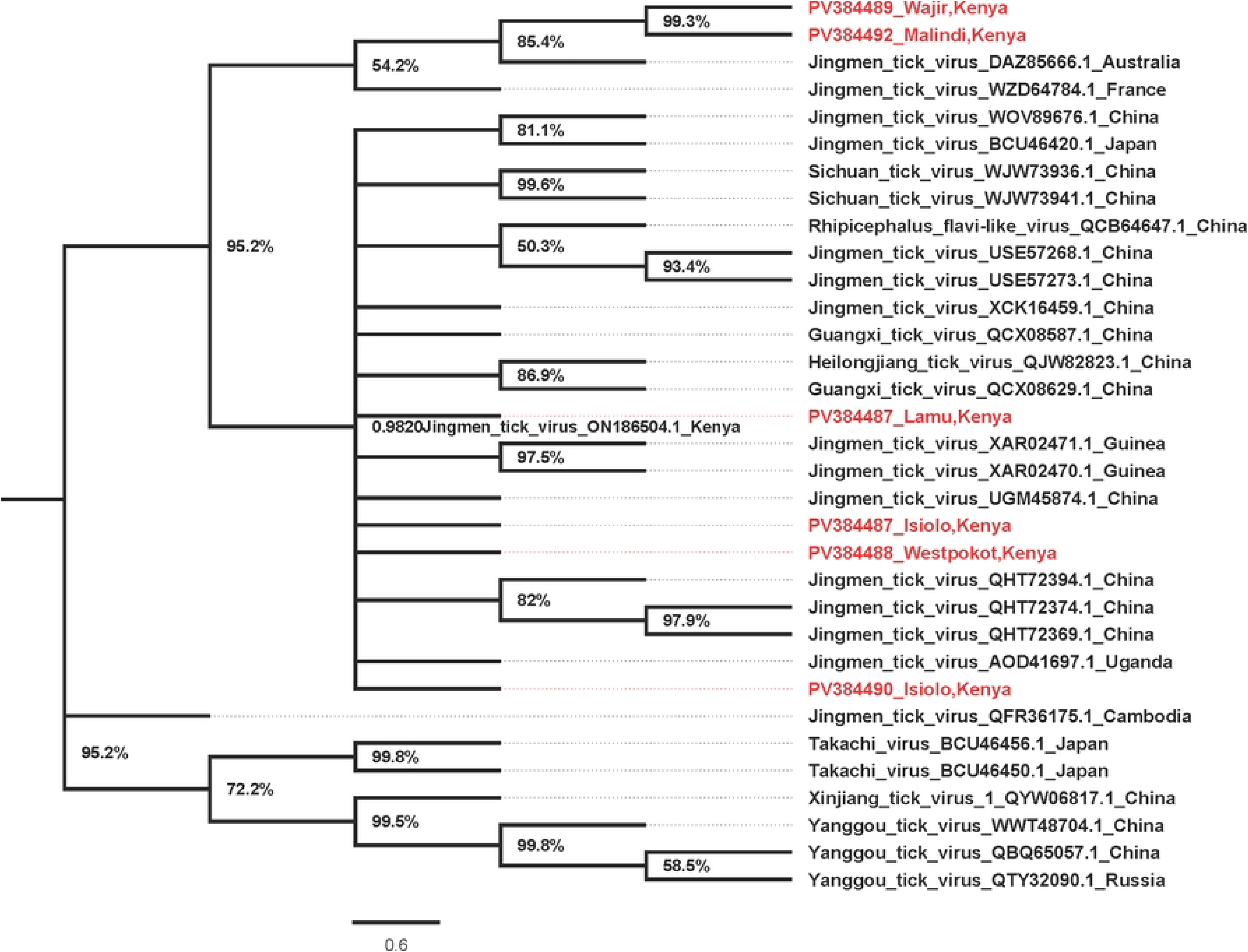
Phylogenetic tree for JMTV segment 1 (NSP1) Amino acid maximum likelihood tree performed in MEGA 7 version 7.0.26 with 1000 bootstrap replications. Tree midpoint rooted and viewed using Figtree version 1.4.4.

**Table 1.** Number of ticks collected across study sites.

|  | Lamu | Isiolo | Kilifi | Wajir | Mandera | West Pokot | Baringo | Total Per Species |
| --- | --- | --- | --- | --- | --- | --- | --- | --- |
| *Amblyomma gemma* | 8 | 133 | 48 |  | 31 |  |  | 220 |
| *Amblyomma lepidium* |  | 12 | 16 | 36 | 1 | 37 |  | 102 |
| *Amblyomma variegatum* |  | 1 | 4 |  | 87 | 34 |  | 126 |
| *Argas persicus* |  |  |  |  |  |  | 300 | 300 |
| *Hyalomma albiparmatum* |  |  | 24 | 7 | 6 |  |  | 37 |
| *Hyalomma anatolicum anatolicum* |  |  |  | 6 |  |  |  | 6 |
| *Hyalomma dromedarii* |  |  | 1 | 42 | 23 |  |  | 66 |
| *Hyalomma impeltatum* |  |  |  | 6 | 1 |  |  | 7 |
| *Hyalomma marginatum rufipes* | 9 | 97 | 47 | 80 | 51 |  |  | 284 |
| *Hyalomma truncatum* |  | 11 | 1 |  |  | 19 | 98 | 129 |
| *Rhipicephalus appendiculatus* |  | 2 | 31 |  | 1 | 34 |  | 68 |
| *Rhipicephalus boophilus microplus* | 6 | 33 | 86 |  |  | 16 | 18 | 159 |
| *Rhipicephalus evertsi evertsi* |  |  | 25 |  | 23 | 5 |  | 53 |
| *Rhipicephalus pulchellus* | 2 | 45 | 8 | 9 | 8 |  |  | 72 |
| **Total Per Site** | 25 | 334 | 291 | 186 | 232 | 145 | 416 | 1629 |

### Prevalence and percent homology of JMTV sequences

The ticks were pooled ≤10 based on tick species, animal host and collection site resulting in 231 pools [29]. Aliquots from the original pools were further pooled into 72 super pools of ≤50 per tick species and site of collection. A total of (38/72) 53% of pools were positive for at least one viral pathogen, with JMTV detected in 87% (33/38) of these pools across all study sites. The Jingmen tick virus segment sequences were deposited in GenBank under the accession numbers PV384450 - PV384535 under BioProject ID: PRJNA1238498. Sequences included in this manuscript (S1 Table).

The study JMTV sequences had a nucleotide homology of 89-94% with JMTV sequences from Kenya, Uganda, China, and Japan, and amino acid homology of 90-99% with JMTV sequences from Kenya, Uganda, China, Laos, and Japan (S2 Table).

### Phylogenetic analyses

JMTV Segment 1 for sequences from Kilifi/Malindi and Wajir clustered together in a well-supported and distinct clade. Those from Lamu, Isiolo, and West Pokot clustered in a second clade together with other strains from around the world (Fig 2). Interestingly, JMTV segment two sequences from Wajir, Kilifi/Malindi, and West Pokot clustered in a similar clade as segment 1. The strains from Russia, France, and Switzerland clustered in a second clade while sequences from Lamu and Isiolo clustered in a third clade with the rest of the strains from around the world (Fig 3). The JMTV segment 3 sequences from West Pokot and Wajir clustered in one clade maintaining the tree topology similar to segment 1 and segment 2 trees.

**Fig 3.**
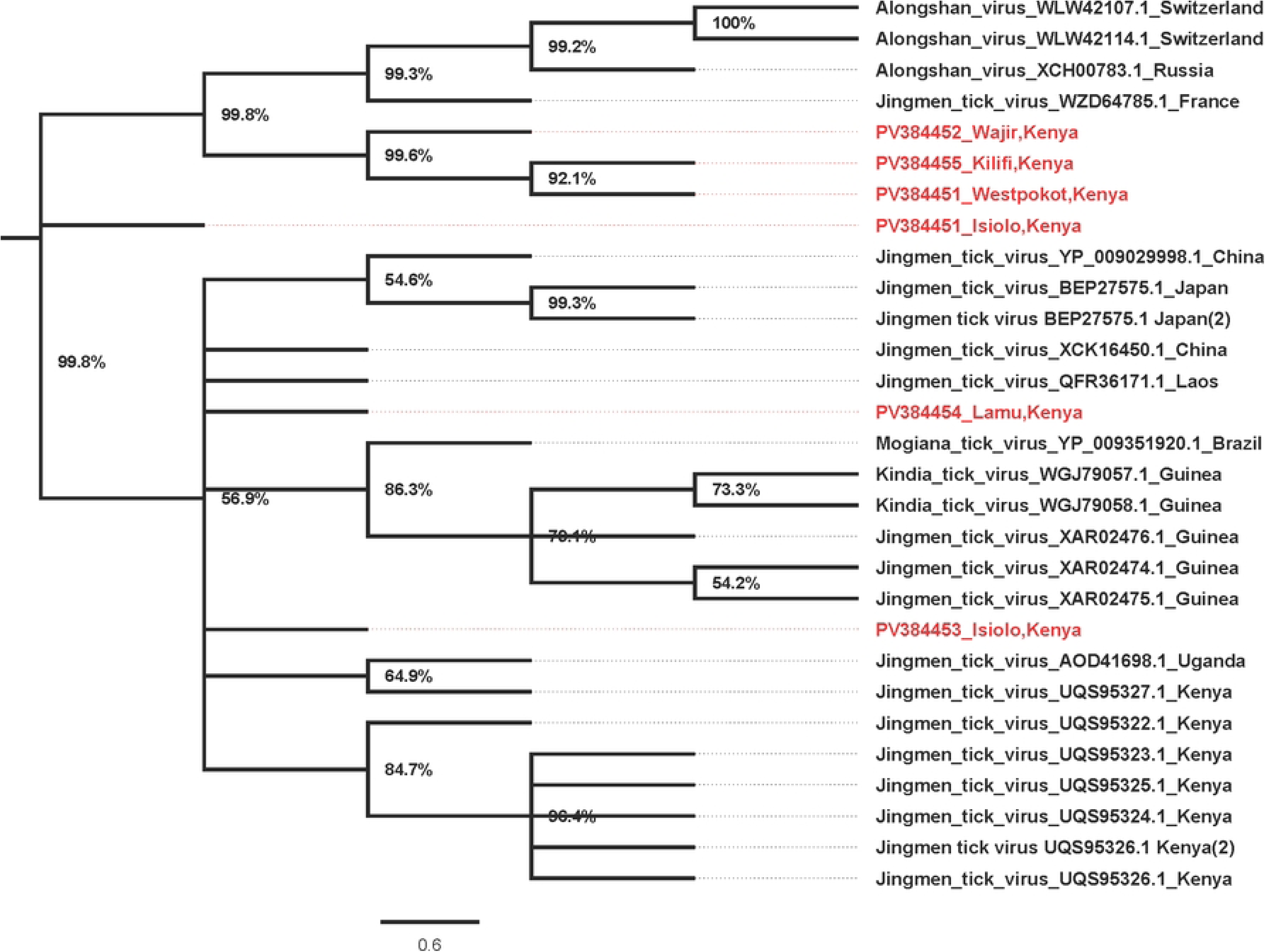
Phylogenetic tree for JMTV segment 2 (VP1) Amino acid maximum likelihood tree performed in MEGA 7 version 7.0.26 with 1000 bootstrap replications. Tree midpoint rooted and viewed using Figtree version 1.4.4.

The segment 3 sequences from Isiolo, Kilifi/Malindi, and Lamu clustered with the rest of the strains from around the world (Fig 4). Finally, for segment 4 JMTV sequences, Wajir, Kilifi/Malindi, and Isiolo clustered in one clade, with Cambodia, Guinea, and China. However, the sequences from Wajir and Cambodia formed distinct branches within the clade. Sequences from West Pokot and Lamu and a second sequence from Isiolo clustered in a second clade with the rest of the strains from around the world clustering together in a third clade (Fig 5).

**Fig 4.**
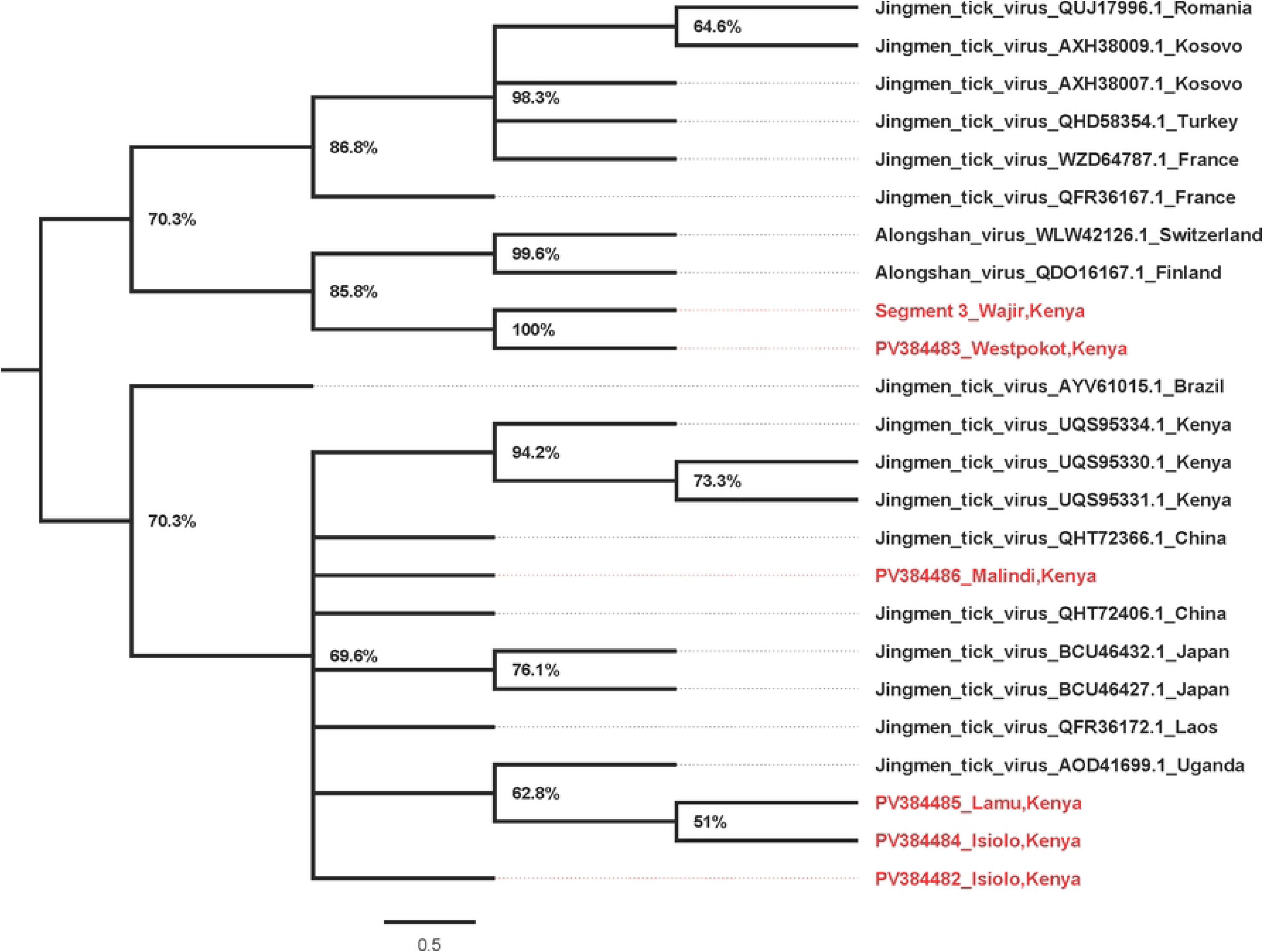
Phylogenetic tree for JMTV segment 3 (NSP2) Amino acid maximum likelihood tree performed in MEGA 7 version 7.0.26 with 1000 bootstrap replications. Tree midpoint rooted and viewed using Figtree version 1.4.4

**Fig 5.**
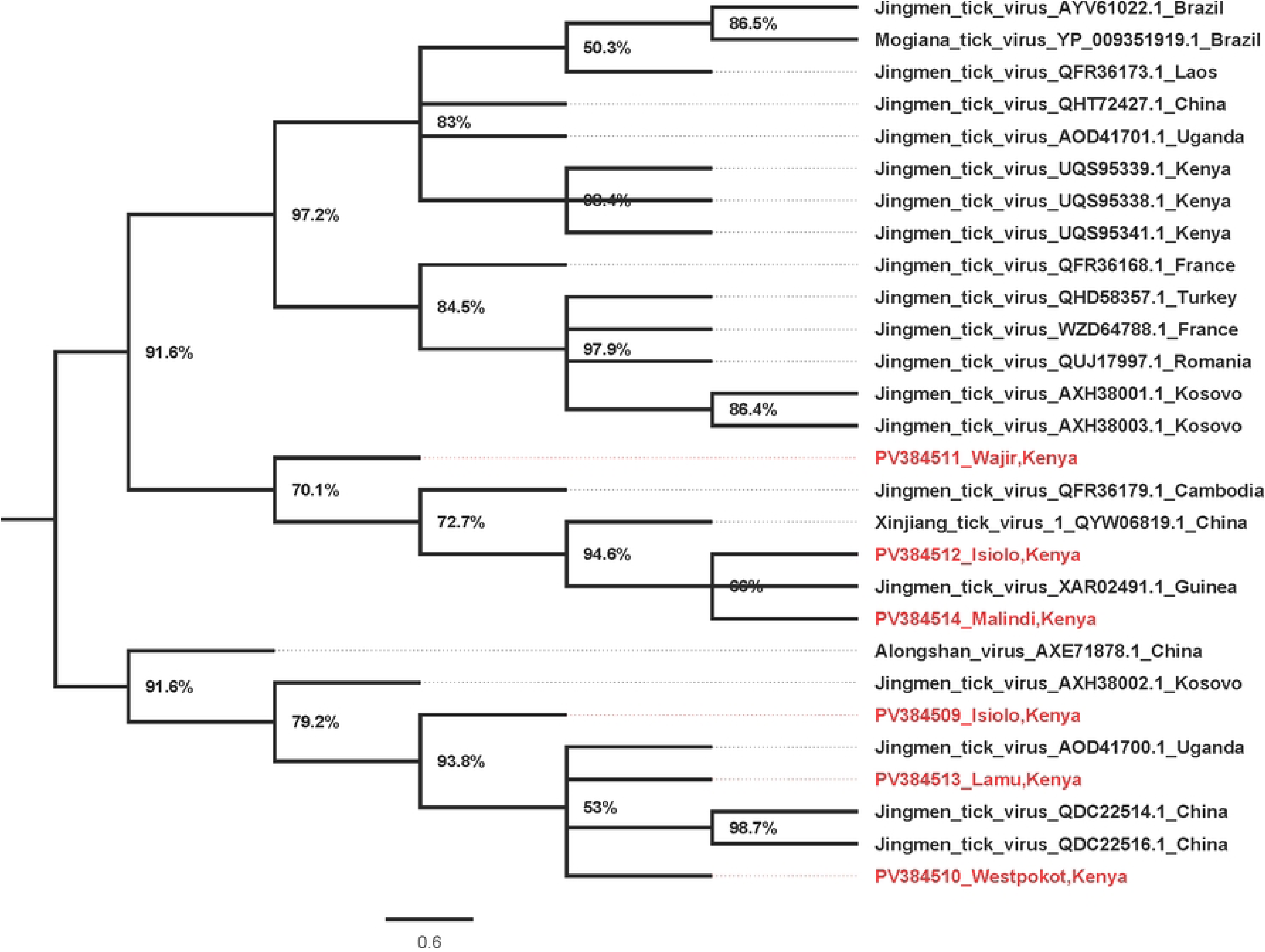
Phylogenetic tree for JMTV segment 4 (VP2&3) Amino acid maximum likelihood tree performed in MEGA 7 version 7.0.26 with 1000 bootstrap replications. Tree midpoint rooted and viewed using Figtree v 1.4.4.

The JMTV time calibrated tree showed one Isiolo branched from ancestral strain in 1950 while the second Isiolo strain, Wajir and Malindi/Kilifi strains branched from their ancestral strain in 2022. The Lamu strain branched from ancestral strain in 1954 while the West Pokot strain branched from ancestral strain in 1943. This suggests varied ancestral origins for JMTVs in the present study, and possible multiple introduction events (Fig 6). The data also shows one of the Isiolo strains, together with those from Wajir and Kilifi/Malindi forming a clade of their own; a second Isiolo strain formed a clade with other JMTV strains from Kenya while the Lamu strain formed a clade with JMTV strain from Uganda and in close proximity to the West Pokot strain (Fig 7). These findings suggest possible inter-country border transmission of JMTV to Kenya from Uganda through the border at West Pokot.

**Fig 6.**
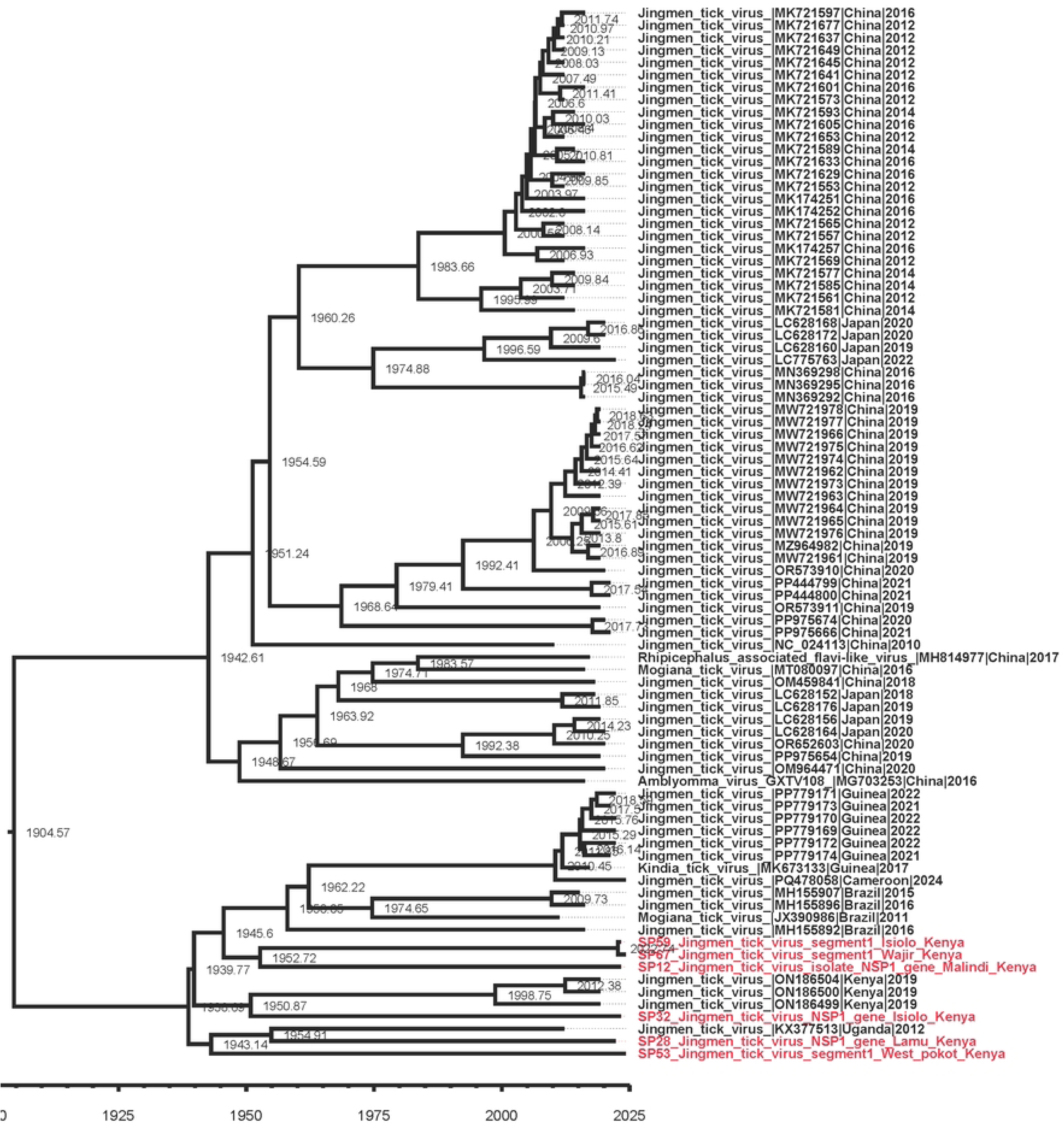
Time calibrated tree showing timescale of JMTV. A phylogenetic tree constructed based on segment 1 (RdRp) of JMTV.

**Fig 7.**
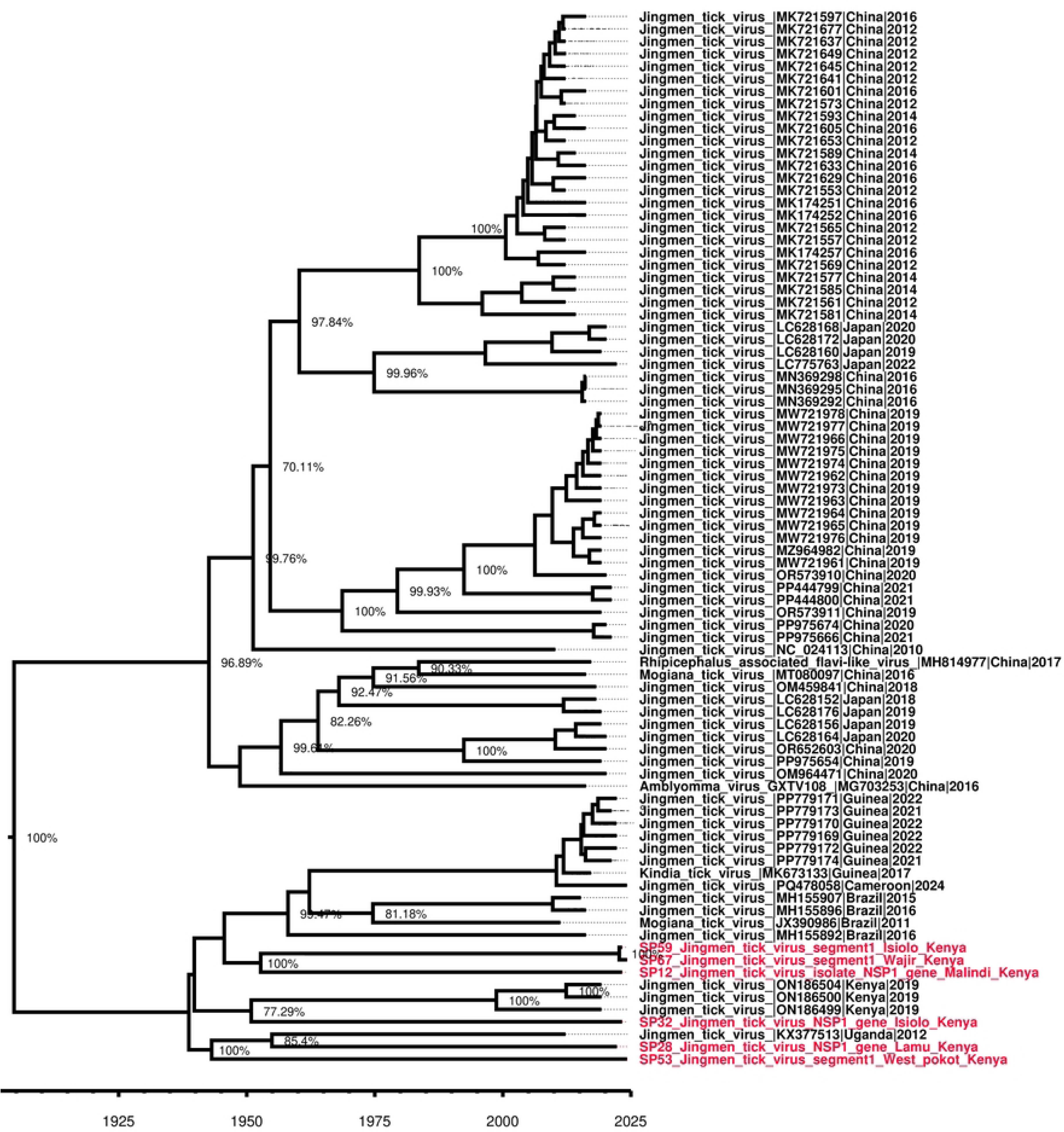
Time calibrated tree showing evolutionary relationships. A phylogenetic tree constructed based on segment 1 (RdRp) of JMTV.

### Natural selection and recombination events

Episodic positive selection analysis using MEME, FUBAR, and BUSTED confirmed selection at codon site 290 in segment 1 and site 30 in segment 4 (Table 2).

**Table 2.**
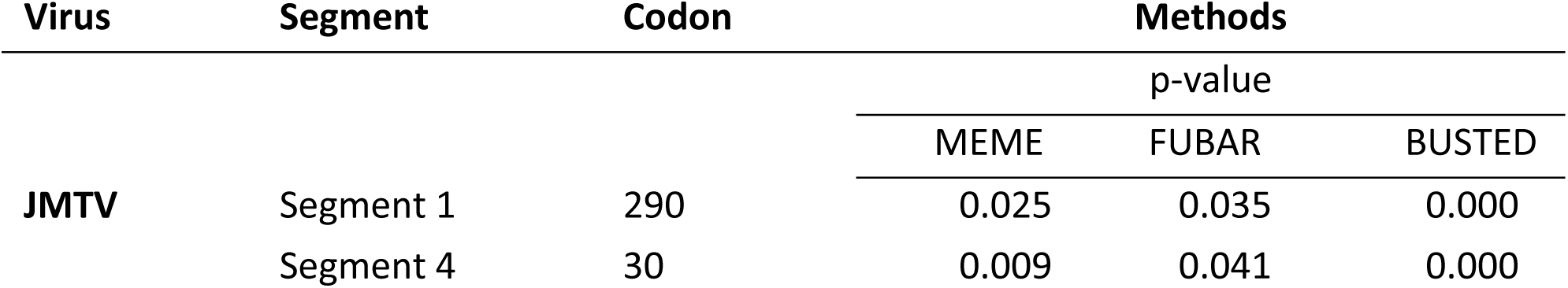
Codon sites under episodic positive selection pressure. Positive selection across JMTV segments using MEME, FUBAR and BUSTED tools of the DataMonkey.

We detected one recombination signal in JMTV segment 2 of the Lamu study strain, PV384454 that was confirmed by 7 of the 8 recombination tools in RDP4 (Table 3), with the Kenyan JMTV strain ON186506 as the parental sequence (S3 Table). The recombination observed in segment 2 is evidenced by BootScan recombination plot (Fig 8).

**Fig 8.**
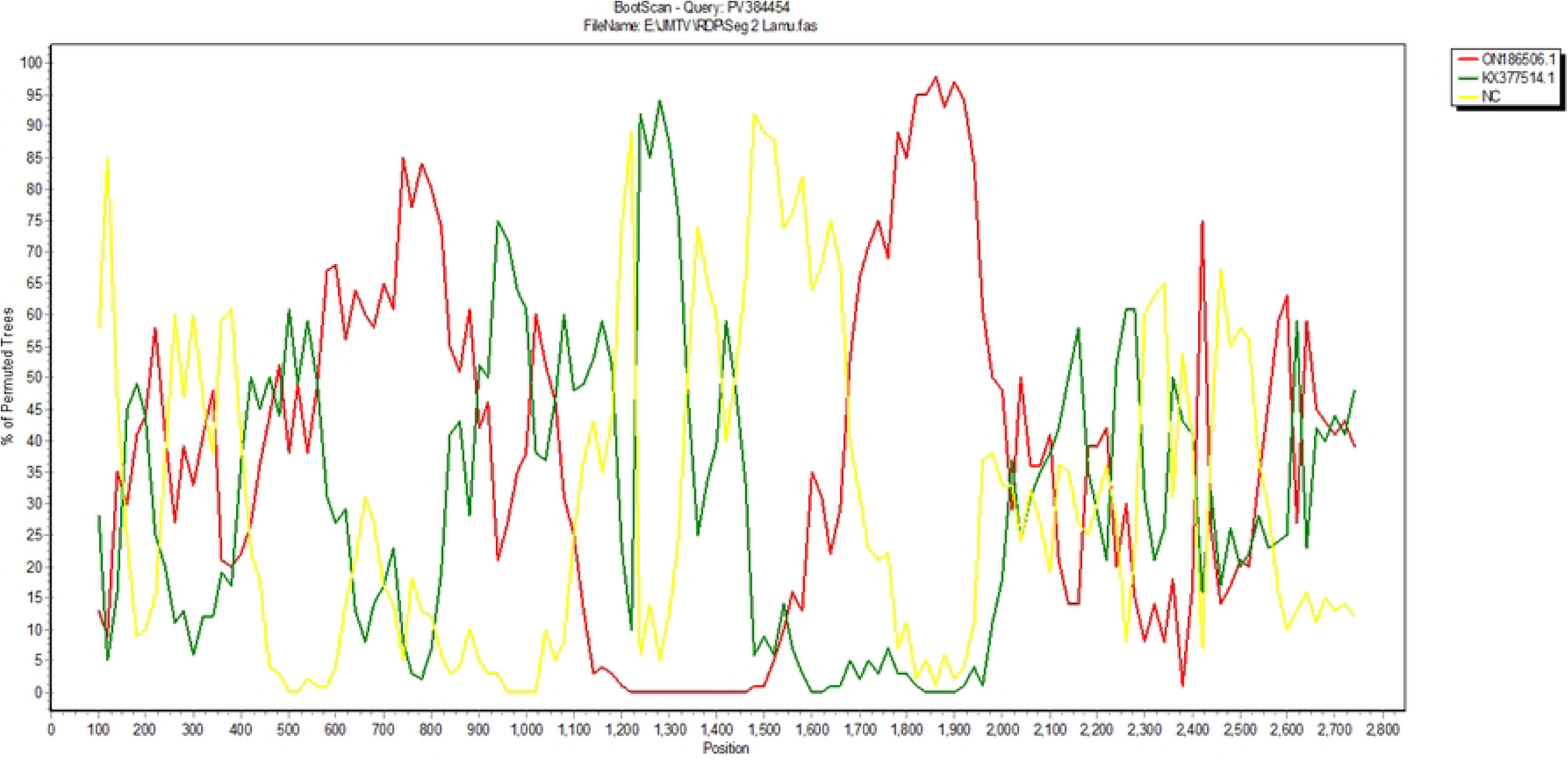
BootScan plot of segment 2 of JMTV genome for strain PV384454, with evidence of recombination. The plot was prepared within a sliding window of 200bp, with a 20bp step size, for 1000 replications (Gap Strip: On, Neighbor - joining, T/t:2.0).

**Table 3.** Recombination signals detected in the JMTV segment 2 using tools in RDP4.

| JMTV | NO | Parental<br>Sequence | Breakpoint<br>positions |  | Detection Methods (P value) |  |  |  |  |  |  |
| --- | --- | --- | --- | --- | --- | --- | --- | --- | --- | --- | --- |
|  |  |  | Begin | End | RDP | GENECONV | BOOTSCAN | MaxChi | Chimera | SiScan | 3Seq |
| segment<br>2 | PV384454 | ON186506.1 | 6 | 124 | 1.129E-27 | 8.040E-50 | - | 7.751E-17 | 6.124E-10 | 4.492E-30 | 1.347E-04 |

## Discussion

Since its initial isolation in China in 2010, the JMTV has been detected widely across the globe in ticks, animal vertebrates, and humans [29]. JMTV was first identified from the *Rhipicephalus boophilus microplus* ticks in China in 2010 [1,2,3,4,6,28,31], however, over time, the virus has also been identified in other tick species [31–33], and in ticks from both wild and domestic animals, highlighting the wide vector and host adaptation of the JMTV virus [28]. RNA viruses are known to evolve through a complex interplay of long-term co-evolution with their hosts and repeated events of host switching [34–36]. Previous JMTV studies have suggested possible genetic reassortment and recombination in JMTV during cross-species transmission [28,36,37]. This rather complex ticks-vertebrates relationship has an impact on evolutionary relationships and consequently increases human infection risk [28]. The current study has obtained partial and near full-length sequences of JMTV strains from *Rhipicephalus boophilus microplus*, *Amblyomma variegatum, Hyalomma dromedarii* and *Amblyomma lepidium* ticks collected from Isiolo, West Pokot, Wajir, Kilifi/Malindi and Lamu regions of Kenya between 2022-2024. The overall positive-rate of JMTV in this study (45.8%) was comparatively higher than observed in a similar JMTV study in Turkey (3.9%) [33]. Comparative prevalence was observed in ticks from other studies in China (53-63%), Brazil (25-67%), Trinidad and Tobago (6-46%) and in the French Antilles (24-77%) [33–35]. In addition, our results are consistent with surveys conducted in all these studies; suggesting an increased risk of JMTV transmission among ticks through co-feeding in animals [28;33]. The geo-phylogenetic distribution of JMTV strains in the current study portrayed diversity in phylogenetic placement with other JMTV strains across the globe; suggesting possible movement between regions [32,36,37]. The time calibrated trees for evolutionary relationships and time scale suggest multiple ancestry origins for the study strains and thus potential varied times of introduction into the country, further explaining the strain divergence across the study sites.

Positive episodic selection on segment 1 and 4 correlate with previous studies that have observed some evolutionary pressures in these JMTV segments [28], despite their conserved nature and important role in function of proteins they encode [28]. The segment 1 of JMTV encodes the non-structural protein NSP1, whose role is to mediate virus replication in the host while segment 4 encodes the viral protein 1 (VP1) and other proteins, related to changes in the virus’s ability to infect or transmit to other hosts. Evolutionary pressures at segments 1 and 4 suggests increased adaptability within the virus and its host for enhanced stability and dominance [34].

A recombination event observed on segment 2 of the Lamu strain PV384454 in comparison to previous strains from Kenya, China, and Uganda, with possible parental sequence to be that of a previous Kenyan strain ON186506 agree with previous studies that found JMTV segment 2 as having the fastest evolutionary rates in comparison to other JMTV segments [29].

## Conclusion

JMTV has been detected not only in tick, but also vertebrate hosts, in both domesticated and wild animals. JMTV has been associated with human disease, both mild and severe; possibly explaining unresolved cases of febrile illnesses. The broad geographic distribution of JMTV and its diverse viral hosts suggest that the JMVs associated with ticks and vertebrates have become globally endemic with future pandemic potential. This study has identified JMTV in 7 pastoral regions of Kenya, genetically characterizing the virus while demonstrating possible ancestral origin and timelines introduction to the country. Future studies to identify the growth potential and rate in relation to flaviviruses will further give more information about the nature of the Jingmen tick virus in Kenya.

## Acknowledgements

We thank Janet Ambale, Nicholas Odemba, Elizabeth Kioko, Jane Thiiru, Solomon Langat and Moshe Dayan for their support in supply procurement and expert contribution to tick sampling and identification. We also thank Samuel Owaka for his contribution to generating the map of sampling sites.

The material has been reviewed by the Walter Reed Army Institute of Research. There is no objection to its presentation and/or publication. The opinions or assertions contained herein are the private views of the authors and are not to be construed as official or as reflecting the true views of the Department of the Army or the Department of Defense.

## Supporting information

**S1 Table. Study JMTV strains and their GenBank Accession numbers.**

**S2 Table. JMTV nucleotide and amino acid percent homology with other global JMTV sequences.**

**S3 Table. JMTV recombination analysis sequences.**

## References

1. Taniguchi S. Detection of Jingmen tick virus in human patient specimens: Emergence of a new tick-borne human disease? EBioMedicine. 2019; 43:18–19. doi: 10.1016/j.ebiom.2019.04.034. PMCID: PMC6558021.

2. Colmant Agathe M, Charrel Rémi, Coutard Bruno. Jingmenviruses: Ubiquitous, understudied, segmented flavi-like viruses. Frontiers in Microbiology. 2022;13. doi:10.3389/fmicb.2022.997058.

3. Maruyama R, Castro-Jorge, Ribeiro C, Gardinassi G, Garcia R, Brandão G, et al. Characterization of divergent flavivirus NS3 and NS5 protein sequences detected in Rhipicephalus microplus ticks from Brazil. Mem Inst Oswaldo Cruz. 2014; 109:38–50. doi: 10.1590/0074-0276130166

4. Qin C, Shi M, Tian H, Lin D, Gao Y, He R, et al. A tick-borne segmented RNA virus contains genome segments derived from unsegmented viral ancestors. PNAS. 2014; 111: 6744–6749. doi: 10.1073/pnas.1324194111

5. Kholodilov S, Litov G, Klimentov S, Belova A, Polienko E, Nikitin A, et al. Isolation and characterization of Alongshan virus in Russia. Viruses. 2020;12: e362. doi: 10.3390/v12040362

6. Ogola O, Anne K, Armanda B, Inga S, Marco M, Dorcus O, et al. Jingmen tick virus in ticks from Kenya. Viruses. 2022;14(5): e1041. doi:10.3390/v14051041

7. Okello-Onen J, Hassan M, Suliman E. Taxonomy of African ticks: an identification manual. International Centre of Insect Physiology and Ecology (ICIPE); 1999.

8. Moulton-Meissner H, Noble-Wang J, Gupta N, Hocevar S, Kallen A, Arduino M. Laboratory replication of filtration procedures associated with Serratia marcescens bloodstream infections in patients receiving compounded amino acid solutions. Am J Health Syst Pharm. 2015;72(15):1285–1291. doi: 10.2146/ajhp150141

9. Alhusein N, Scott J, Kasprzyk-Hordern B, Bolhuis A. Development of a filter to prevent infections with spore-forming bacteria in injecting drug users. Harm Reduct J. 2016;13(1):33. doi: 10.1186/s12954-016-0122-1

10. Perkel M. Terra takes the pain out of ‘omics’ computing in the cloud. Nature. 2022; 601(7891), 154–155. doi: 10.1038/d41586-021-03822-7

11. Anthony B, Marc L, Bjoern U. Trimmomatic: a flexible trimmer for Illumina sequence data. 2014. Bioinformatic;30(15): 2114–2120. doi: 10.1093/bioinformatics/btu170

12. Chen S. Ultrafast one-pass FASTQ data preprocessing, quality control, and deduplication using fastp. Imeta. 2023;2(2): e107. doi: 10.1002/imt2.107

13. Wood E, Lu J, Langmead B. Improved metagenomic analysis with Kraken 2. Genome biology. 2019; 20(1), 257. doi: 10.1186/s13059-019-1891-0

14. Nurk S, Meleshko D, Korobeynikov A, Pevzner PA. metaSPAdes: a new versatile metagenomic assembler. Genome Res. 2017;27(5):824–834. doi: 10.1101/gr.213959.116

15. Petr D, James B, Jennifer L, John M, Valeriu O, Martin P, et al. Twelve years of SAMtools and BCFtools. GigaScience.2021; 10 (2): e008. doi:10.1093/gigascience/giab008

16. Heng Li. Minimap2: pairwise alignment for nucleotide sequences. Bioinformatics. 2018; 34(18): 3094–3100. doi:10.1093/bioinformatics/bty191

17. Walker BJ, Abeel T, Shea T, Priest M, Abouelliel A, Sakthikumar S, et al. Pilon: an integrated tool for comprehensive microbial variant detection and genome assembly improvement. PLoS One. 2014; 9(11): e112963. doi: 10.1371/journal.pone.0112963

18. Gurevich A, Saveliev V, Vyahhi N, Tesler G. QUAST: quality assessment tool for genome assemblies. Bioinformatics. 2013;29(8):1072–1075. doi: 10.1093/bioinformatics/btt086

19. Kumar S, Stecher G, Tamura K. MEGA7: Molecular evolutionary genetics analysis version 7.0 for bigger datasets. Mol Biol Evol. 2016;33(7):1870-1874. doi: 10.1093/molbev/msw054

20. Stéphane G, Jean-François D, Vincent L, Maria A, Wim H, Olivier G. New algorithms and methods to estimate maximum-likelihood phylogenies: assessing the performance of PhyML 3.0. Systematic Biology.2010; 59 (3): 307–321. doi:10.1093/sysbio/syq010

21. Rambaut A. Figtree ver 1.4.4. Institute of Evolutionary Biology. University of Edinburgh, Edinburgh 2018.

22. Kosakovsky Pond SL, Frost SD. Not so different after all: a comparison of methods for detecting amino acid sites under selection. Mol Biol Evol. 2005;22(5):1208–1222. doi: 10.1093/molbev/msi105

23. Murrell B, Wertheim JO, Moola S, Weighill T, Scheffler K, Kosakovsky Pond SL. Detecting individual sites subject to episodic diversifying selection. PLoS Genet. 2012;8(7): e1002764. doi: 10.1371/journal.pgen.1002764

24. Ben M, Sasha M, Amandla M, Thomas W, Daniel S, Sergei L., et al. FUBAR: A Fast, Unconstrained Bayesian AppRoximation for inferring selection. Molecular Biology and Evolution. 2013; 30, (5):1196–1205. doi:10.1093/molbev/mst030

25. Ben M, Steven W, Martin DS, Joel OW, Sasha M, Anthony A, et al. Gene-wide identification of episodic selection. Molecular Biology and Evolution. 2015;32(5):1365–1371. doi:10.1093/molbev/msv035

26. Darren PM, Ben M, Michael G, Arjun K, Brejnev M. RDP4: Detection and analysis of recombination patterns in virus genomes. Virus Evolution. 2015;1(1). doi:10.1093/ve/vev003

27. Jaya FR, Brito BP, Darling AE. Evaluation of recombination detection methods for viral sequencing. Virus Evolution. 2023;9(2). doi: 10.1093/ve/vead066.

28. Weiyi L, Rongting L, Xiaomin T, Jinzhi C, Lin Z, Zhengling S, et al. Genomics evolution of Jingmen viruses associated with ticks and vertebrates. Genomics. 2023;115(6). doi: 10.1016/j.ygeno.2023.110734

29. Lole KS, Bollinger RC, Paranjape RS, Gadkari D, Kulkarni SS, Novak NG, et al. Full-length human immunodeficiency virus type 1 genomes from subtype C-infected sero converters in India, with evidence of inter subtype recombination. Journal of virology. 1999);73(1):152–160. doi: 10.1128/JVI.73.1.152-160.1999

30. Edouard de Castro, Christian AS, Alexandre G, Virginie B, Petra S. L, Elisabeth G, et al. ScanProsite: detection of PROSITE signature matches and ProRule-associated functional and structural residues in proteins. Nucleic Acids Research. 2006; 34 (2):362–365. doi: 10.1093/nar/gkl124

31. Dinçer E, Hacıoğlu S, Kar S, Emanet N, Brinkmann A, Nitsche A, et al. Survey and characterization of Jingmen tick virus variants. Viruses. 2019;11(11):1071. doi: 10.3390/v11111071

32. Ergunay K, Mutinda M, Bourke B, Justi SA, Caicedo-Quiroga L, Kamau J, et al. Metagenomic investigation of ticks from Kenyan wildlife reveals diverse microbial pathogens and new country pathogen records. Frontiers in Microbiology.2022;13. doi: 10.3389/fmicb.2022.932224

33. Zhang Y, Li Z, Pang Z, Wu Z, Lin Z, Niu G. Identification of Jingmen tick virus (JMTV) in Amblyomma testudinarium from Fujian Province, southeastern China. Parasit Vectors. 2022;15(1):339. doi: 10.1186/s13071-022-05478-2

34. Shi M, Lin XD, Tian JH, Chen LJ, Chen X, Li CX, et al. Redefining the invertebrate RNA virosphere. Nature. 2016;540(7634):539-543. doi: 10.1038/nature20167

35. Li CX, Shi M, Tian JH, Lin XD, Kang YJ, Chen LJ, et al. Unprecedented genomic diversity of RNA viruses in arthropods reveals the ancestry of negative-sense RNA viruses. Elife. 2015;4:e05378. doi: 10.7554/eLife.05378

36. Wu Z, Zhang M, Zhang Y, Lu K, Zhu W, Feng S, et al. Jingmen tick virus: an emerging arbovirus with a global threat. mSphere. 2023;8(5): e0028123. doi: 10.1128/msphere.00281-23.

37. Kiwan P, Lopez E, Gasparine M, Piorkowski G, Colmant A, Paguem A, et al. First detection and molecular characterization of Jingmen tick virus with a high occurrence in Rhipicephalus (Boophilus) microplus collected from livestock in Cameroon (2024). Parasit Vectors. 2025;18(1):41. doi: 10.1186/s13071-025-06670-w

